# Predicting single-cell responses to novel genetic perturbations with optimal transport

**DOI:** 10.64898/2026.01.08.698172

**Authors:** Chau Do, Harri Lähdesmäki

## Abstract

The combination of pooled genetic perturbation screening with single-cell RNA-sequencing has allowed for large-scale profiling of single-cell transcriptional responses, providing valuable insights into gene function and cellular behavior. There is a need for computational methods capable of predicting single-cell responses to *novel* genetic perturbations, thereby allowing for the efficient exploration of the vast combinatorial space of potential multi-gene perturbations. Optimal transport (OT) presents a compelling framework to model this type of data given the lack of true control-perturbed cell pairings. However, existing neural OT models are generally constrained to the perturbations already seen in training, while the use of OT theory in methods capable of modeling novel genetic perturbations remains limited. To bridge this gap, we present an OT-based method to predict single-cell transcriptional responses to novel genetic perturbations. Leveraging a flexible attention-based aggregation approach, our method integrates gene representations from multiple sources of prior knowledge, spanning from functional descriptions of genes to gene-gene relationships, producing a biologically informed representation of the perturbation. Moreover, we utilize an OT-based loss to align predicted and observed perturbed cell populations, avoiding the need to assign random control-perturbed cell pairings. Experiments on three benchmark datasets demonstrate the highly competitive performance of our model compared to state-of-the-art methods. Further evaluations on differential expression analysis and genetic interaction modeling demonstrate the biological relevance and potential utility of our model across diverse applications.

## 1. Introduction

Large-scale genetic perturbation experiments provide valuable insights into how genomic components interact and regulate cellular behaviors. The development of Perturb-seq technology, which combines pooled CRISPR-based genetic perturbations and scRNA-seq, has enabled large-scale profiling of single-cell transcriptional responses. Perturb-seq has promising applications in cell engineering and targeted therapy [1]. Despite such advancements, the combinatorial space of possible multigene combinations remains expensive and labor-intensive to explore through experimental perturbations. This raises the need for computational tools that are capable of modeling perturbation outcomes *in silico*, allowing for the efficient navigation and exploration of the genome-scale perturbation space.

In recent years, several tools have been developed to predict the transcriptional outcomes of genetic perturbations. ScGEN [2] employs a variational autoencoder architecture and models the effect of genetic perturbations as simple vector additions in the latent space. CPA [3] also uses an autoencoder framework and is additionally capable of modeling the combined effect of multiple perturbations. On the other hand, CellOT [4] employs neural optimal transport (OT) to map cells from the control population to the perturbed population.

However, these tools are not capable of predicting the outcomes of novel genetic perturbations, which involve genes not previously seen perturbed during training. To achieve this, it is essential to be able to meaningfully represent the unobserved genes in the same latent space as the genes perturbed in the training set. For instance, Roohani et al. [1] introduced GEARS, a graph neural network (GNN)-based approach that leverages a knowledge graph of gene relationships to embed the unobserved genes. TxPert [5] extends this idea to multiple knowledge graphs to leverage different sources of gene relationship information, while AttentionPert [6] integrates GNNs with multi-head attention to aggregate the perturbation effects of multiple genes. Instead of relying on knowledge graphs, GenePert [7] and scLAMBDA [8] utilize large-language models (LLMs) to embed the unobserved genes with their text-based descriptions.

A further challenge is that scRNA-seq procedures typically destroy cells, so the control and perturbed cells are unpaired, meaning it is not possible to observe the expression of a single cell both before and after perturbation. This complicates analysis due to the inherent heterogeneity of single cells. To deal with the lack of a true control during training, some models (scGEN, GEARS, TxPert, AttentionPert) either pair up random, possibly batch-matched control and perturbed cells, or use a pseudobulk-averaged control profile. This strategy captures the average perturbation effect yet may not sufficiently account for cellular heterogeneity. Other models (CPA, scLAMBDA) employ a latent disentanglement approach—during training, the model inputs the expressions of perturbed cells, disentangles the perturbed expressions into basal state representations and perturbation effects in the latent space, and finally reconstructs the perturbed expressions as output. While this approach does not make any assumptions about control-perturbed cell pairings, it relies on complete disentanglement between the basal state and perturbation effects, and incomplete disentanglement may lead to suboptimal performance.

A promising alternative is to use OT to directly construct a mapping of cells from the control population to the perturbed population. However, existing OT-based perturbation modeling tools, such as CellOT, are generally constrained to perturbations previously observed during training and do not support generalization to unseen perturbations.

To bridge this gap, we introduce a deep generative model for the prediction of single-cell transcriptomic responses to novel genetic perturbations. With flexible modeling capacity, our approach integrates biological knowledge from multiple sources, including text-based gene descriptions, Gene Ontology (GO) enrichment, and gene functional associations, to generate biologically informed representations of perturbed genes. The contribution of each source of prior knowledge to the gene representations, as well as the contribution of each gene to the multi-gene perturbation representations, are dynamically weighted with an attention-based aggregation function. Moreover, during training, our model utilizes an OT-based loss to align the distribution of the predicted perturbed cells with that of the ground truth, effectively solving the problem of unpaired perturbed-control cells while preserving cellular heterogeneity. This single-cell loss is further combined with a pseudobulk, averaged expression loss, which encourages the predicted perturbed cells to match the ground truth at both the population level and the cell level.

We benchmark our model against three state-of-the-art models on both single-gene and multi-gene perturbation outcome prediction tasks. Across three Perturb-seq datasets and diverse evaluation metrics, our model consistently demonstrates competitive performance. Moreover, ablation test results show that incorporating multiple sources of prior knowledge and utilizing an OT-based loss generally improve performance, validating the effectiveness of such design choices. Additional evaluations on genetic interaction (GI) modeling and differential expression (DE) analysis tasks further demonstrate the utility of our model across diverse biological applications.

## 2. Methods

### 2.1. Overview

Given the gene expressions of control/unperturbed cells and the representations of the perturbed genes (possibly from multiple knowledge sources), our model generates the expressions of cells post-perturbation. Adapting the architecture proposed by Qi et al. [9], our model includes an encoder, a decoder, and multiple gene adapters, each corresponding to gene representations from a source of prior knowledge. For each gene, representations from different sources are embedded via their corresponding adapters, then weighted and combined using an attention-based aggregation function. This generates one embedding vector for each gene. If the perturbation involves multiple genes, gene embeddings are again weighted and aggregated to produce the perturbation representation. The control expressions are then encoded and probabilistically decoded conditioned on the perturbation representation, to generate the perturbed expressions. Training proceeds in two phases: the pre-training phase where the model learns to reconstruct the expression profiles of control and perturbed cells, and the main training phase where the model learns to map the control to the perturbed cell population. The predicted and ground-truth perturbed expression profiles are aligned using a combination of 2-Wasserstein loss across all cells and a mean squared error (MSE) loss on pseudobulk-averaged expression vectors. An overview of our model is given in Fig. 1.

**Fig. 1.**
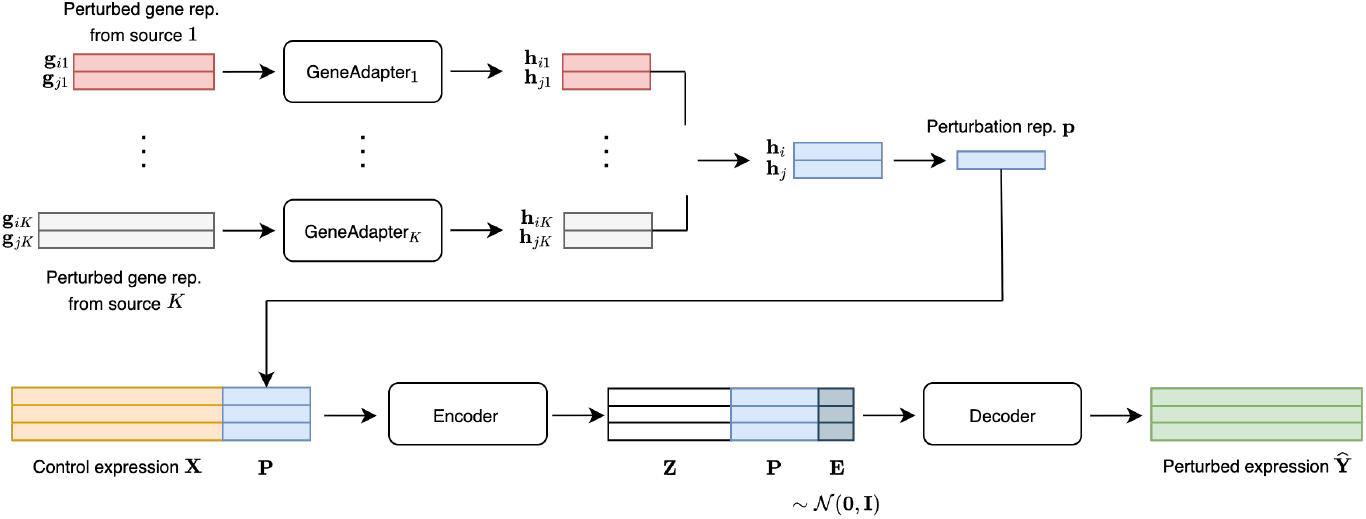
Workflow of the proposed model, demonstrated on two perturbed genes *i* and *j* with representations from sources 1, …, *K*.

### 2.2. Model architecture

The expression profiles of control cells are denoted as 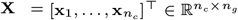, where 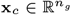 is the expression vector of cell *c, n*_*c*_ is the number of control cells, and *n*_*g*_ is the number of genes. We assume *P* perturbation experiments. For perturbation *p* ∈ {1, …, *P*}, let ℐ_*p*_ ⊆ {1, …, *n*_*g*_} denote the indices of the perturbed genes. The post-perturbation expression profiles of cells in perturbation *p* are denoted as 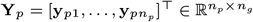, where 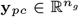, and *n*_*p*_ is the number of cells with perturbation *p*. Assume there are *K* sources of gene representations available for each perturbed gene *i* ∈ {1, …, *n*_*g*_}, collectively denoted as a list **G**_*i*_ = (**g**_*i*1_, …, **g**_*iK*_), where 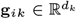 denotes the *d*_*k*_-dimensional representation of gene *i* from source *k*. Given the control cell profiles **X** and gene representations **G**_*i*_ for all genes *i* ∈ ℐ_*p*_ in perturbation *p*, our model generates the predicted perturbed expression profiles 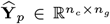. For clarity of notation, we drop the perturbation index *p* and present our method for a single perturbation experiment.

#### 2.2.1. Perturbation adapters

Each perturbed gene representation 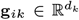 is first mapped to a lower-dimensional gene representation 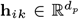 using the gene adapter for knowledge source *k*:

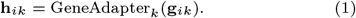

Here, GeneAdapter_*k*_(·) is modeled as a multi-layer perceptron (MLP) with knowledge source-specific learnable parameters.

Next, we aggregate the representations **h**_*ik*_ across knowledge sources *k* using an attention-based aggregation function to produce a single representation 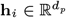 for each gene *i*

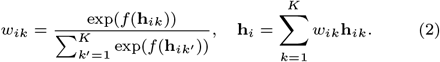

Here, *w*_*ik*_ ∈ ℝ denotes the importance weight of source *k* for gene *i*, and *f* is an MLP with learnable parameters.

In the case of single-gene perturbation (|ℐ| = 1), the perturbation representation 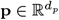 is the representation of its only perturbed gene *i*, i.e., **p** = **h**_*i*_. In the case of multi-gene perturbation, **p** is derived by weighting and aggregating its perturbed genes:

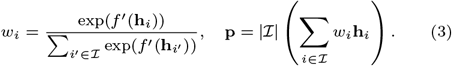

Here, *w*_*i*_ ∈ ℝ denotes the importance weight of gene *i* in the multi-gene perturbation, and *f* ^′^ is modeled as an MLP with learnable parameters. Note that Eq. 3 differs from Eq. 2 in that the aggregated gene representation is scaled by the number of perturbed genes |ℐ| to produce the perturbation representation **p**. This reflects the intuition that the effect of the perturbation is more pronounced when more genes are involved.

#### 2.2.2. Encoder and decoder

Conditioned on the perturbation representation **p**, the control expression profiles 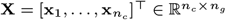 are mapped to the latent representations 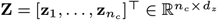 via the encoder. The encoder operates on each cell (i.e., row of the matrix) separately, and the encoding of the entire cell population is collectively denoted as

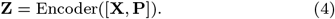

The perturbation matrix 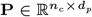 is obtained by repeating the perturbation representation **p** *n*_*c*_ times along the first dimension. Formally, 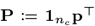, where 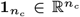 is a vector of ones. The encoder is modeled as an MLP with learnable parameters.

Finally, the latent representations **Z** are probabilistically decoded, conditioned on **p**, to generate the predicted perturbed expression profiles 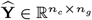:

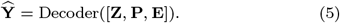

Here, 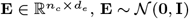 is the noise matrix. Similarly to the encoding step, the decoder maps each cell (i.e., row of the matrix) separately. Note that instead of explicitly defining a likelihood model for 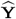, we use the push-forward technique to map **Z** and **P** together with the simple Gaussian base distribution to the post-perturbation cell samples. The decoder is modeled as an MLP with learnable parameters.

### 2.3. Model training

Training consists of the pre-training and main training phases.

#### 2.3.1. Pre-training phase

During the pre-training phase, the model is trained to reconstruct the control expressions **X** and the perturbed expressions **Y**. Since there is no perturbation applied, the perturbation representation **P** is replaced with 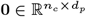. Specifically, for reconstructing **X** and **Y**:

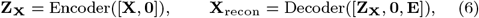

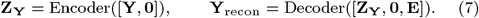

The training objective is the MSE between **X, Y** and their reconstructions **X**_recon_, **Y**_recon_.

#### 2.3.2. Main training phase

During the main training phase, the model is trained to generate the perturbed cells given the control cells and the representations of perturbed genes, as described in Sec. 2.2. The training objective ℒ balances three loss terms: the single-cell loss ℒ_sc_, the pseudobulk loss ℒ_pb_, and the reconstruction loss ℒ_recon_.

ℒ_sc_ is the quadratic 2-Wasserstein distance between the predicted perturbed cell population 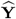 and the ground truth **Y**, with higher weights assigned to the top 20 differentially expressed genes (DEGs) of the perturbation:

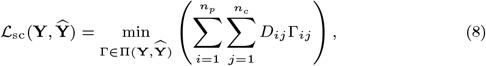

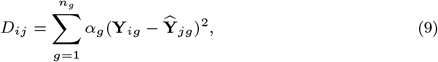

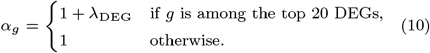

Here, **Y** and 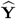 are treated as empirical distributions, with all cells (points) given equal mass. 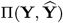 is the set of all possible transport plans from **Y** to 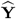. For 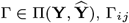 indicates how much mass to transport from point *i* in **Y** to point *j* in 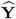. *D*_*ij*_ is the cost of transporting one unit of mass from point *i* in **Y** to point *j* in 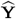, which is the squared Euclidean distance between the two points, but with higher weights *α*_*g*_ assigned to dimensions corresponding to the top 20 DEGs. ℒ_sc_ is approximated with the Sinkhorn divergence [10]. ℒ_pb_ is defined as

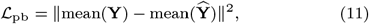

where mean(·) averages the expression profiles across cells to produce the pseudobulk expression vector. The third loss term, ℒ_recon_ is the loss used in the pre-training phase, which is the MSE between **X, Y** and their reconstructions **X**_recon_, **Y**_recon_, as described in Sec. 2.3.1.

The final training loss is defined as

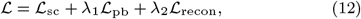

where *λ*_1_ and *λ*_2_ are weights to balance the loss terms. The pre-training and the main training phases are implemented using data from all training perturbations. Conceptually, this loss function guides the model to generate predictions that align with the ground truth, both population-wise via the Wasserstein loss and on average via the pseudobulk L2 loss. In addition, the reconstruction term encourages the model to comply with the intuition that the expression profiles of cells should remain unchanged when no perturbation is applied. Further details on model training can be found in Supplementary Sec. S2.

## 3. Experiments

### 3.1. Experiment setup

#### 3.1.1. Datasets

We benchmark the performance of our model on three commonly-used Perturb-seq datasets: Adamson et al. [11], Replogle et al. [12], and Norman et al. [13]. The Adamson et al. dataset contains single-gene perturbations from a CRISPR interference (CRISPRi) experiment on K562 human erythroleukemic cells. The Replogle et al. dataset also includes single-gene perturbations from a genome-scale CRISPRi screen on retinal pigment epithelial (RPE1) cells. The Norman et al. dataset is a CRISPR activation (CRISPRa) dataset with both single-gene and double-gene perturbations on K562 cells.

The three datasets are preprocessed by Roohani et al. [1]. Preprocessing includes quality control, normalization of total counts per cell, and log-transformation of the normalized counts. Each dataset is further filtered to the top 5000 highly variable genes (HVGs). For the Norman et al. and the Adamson et al. datasets, perturbed genes are additionally included in the expression profiles if they are not already among the top 5000 HVGs. For each perturbation, the top 20 DEGs relative to the control expression profiles are also identified. Moreover, since our model and all competing models rely on external databases or specific strategies to obtain information on perturbed genes, some perturbations could not be predicted as the genes involved are missing from these sources. For consistency and fairness, perturbations that could not be predicted either by our model or by competing models are removed from each dataset, and all models are evaluated on the same set of perturbations. After preprocessing, the Adamson et al. dataset contains 81 single-gene perturbations, the Replogle et al. dataset contains 645 single-gene perturbations, and the Norman et al. dataset contains 102 single-gene perturbations and 128 double-gene perturbations. Further details on train-test splitting in each experiment can be found in Supplementary Sec. S2.

#### 3.1.2. Baselines

We benchmark the performance of our model against three existing methods for perturbation response prediction: GEARS [1], scLAMBDA [8], and scGPT [14]. GEARS is a GNN-based method that leverages a network of pathway-related GO similarity between genes to derive embeddings for novel perturbed genes [15, 16]. ScLAMBDA is a deep generative model that employs a latent disentanglement approach and utilizes LLM embeddings of gene descriptions from NCBI (i.e., GenePT embeddings) to predict responses to novel perturbations [17, 18]. ScGPT is a foundation model for single-cell multi-omics, leveraging a transformer-based architecture. As it inputs a vector indicating the perturbation condition (e.g., perturbed or not perturbed) of each gene to derive the final cell embedding, it requires that perturbed genes have their expression values included in the dataset. For additional information about the baseline models, please see Supplementary Sec. S7.

#### 3.1.3. External sources of gene representations

To represent perturbed genes, we utilize information from three external sources (*K* = 3): LLM embeddings of gene descriptions from NCBI [17, 18], gene similarity based on pathway-related GO terms [15, 16], and functional gene associations from STRING [19]. The LLM embeddings are 3072-dimensional vectors (*d*_1_ = 3072) generated using the GPT-3.5 model, following the approach of scLAMBDA [8]. The GO representation of each perturbed gene contains *n*_*g*_ elements (*d*_2_ = *n*_*g*_), where *n*_*g*_ denotes the number of genes included in the expression profiles. The *i*^th^ element of this vector represents the Jaccard index between the two sets of pathway-related GO terms associated with the perturbed gene and the *i*^th^ gene in the expression profiles. Similarly, the STRING feature vector also contains *n*_*g*_ elements (*d*_3_ = *n*_*g*_), with the *i*^th^ element corresponding to the STRING association score between the perturbed gene and the *i*^th^ gene in the expression profiles. Combining these three sources of prior knowledge generates a biologically informed representation that integrates information on the characteristics, associations, and biological role of each perturbed gene.

#### 3.1.4. Evaluation metrics

To evaluate the alignment between the predicted perturbed cells and the ground truth, we consider four metrics that are commonly used to evaluate perturbation outcome prediction: delta Pearson’s correlation coefficient (ΔPCC) on all genes, the 2-Wasserstein (W2) distance on all genes, and ΔPCC and W2 distance on the top 20 DEGs (ΔPCC-DEG and W2-DEG). ΔPCC is defined as

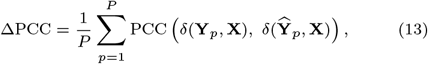

where *δ*(**Y, X**) = mean(**Y**) − mean(**X**) is the pseudobulk expression change, PCC(·) is the PCC function, *P* is the number of test perturbations, **Y**_*p*_ and 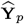 are the ground truth and predicted perturbed cells of perturbation *p*. The W2 distance is calculated on **Y**_*p*_ and 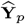 and averaged across test perturbations. ΔPCC and W2 distance on the top 20 DEGs are defined similarly, except that **Y**_*p*_ and 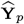 are limited to the top 20 DEGs of perturbation *p*. These four metrics assess both the pseudobulk and single-cell alignment between **Y**_*p*_ and 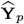 with an emphasis on top DEGs, which are often biologically relevant genes.

### 3.2. Perturbation response prediction

We first evaluate the performance of our model and competing models on the perturbation response prediction task. For each dataset, we randomly split the perturbations into training, validation, and test sets as described in Supplementary Sec. S2, and compute the four evaluation metrics described in Sec. 3.1.4 on the test perturbations. The ΔPCC and W2 distance of all models on the Norman et al. dataset over 10 random data splits are given in Fig. 2. Results on other datasets and metrics are given in Supplementary Figs. S1, S2, and S3.

**Fig. 2.**
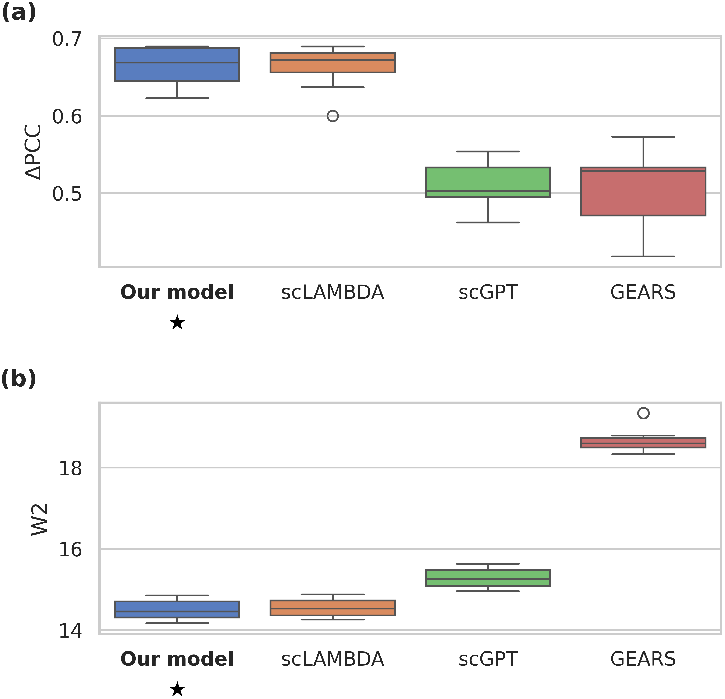
Model performance on the Norman et al. dataset, evaluated using (a) Δ PCC and (b) W2 distance. The model with the best average result on each metric is bolded and starred.

Overall, our model performs competitively with scLAMBDA and outperforms both GEARS and scGPT across all datasets and evaluation metrics. Specifically, it achieves the best average performance across all four metrics on the Norman et al. dataset, and across three out of four metrics on the other two datasets. Even though the performance gain of our model over scLAMBDA is moderate in magnitude, it achieves statistically significant reductions in the W2 distance (*p <* 0.05, one-sided paired *t*-test) on all three datasets, indicating a consistent improvement in the alignment between the predicted and ground truth cell distributions.

### 3.3. Ablation test

To assess the contributions of the core components in our model’s design, we perform an ablation test. First, we evaluate the effectiveness of incorporating multiple knowledge sources by removing the gene representations one by one. Specifically, we first remove the STRING representations, leaving the LLM and GO representations. We subsequently remove the GO representations, leaving only the LLM representations, which are those used by scLAMBDA. The resulting ablated models are referred to as the LLM + GO model and the LLM model, respectively. Moreover, we assess the impact of using an OT-based loss by replacing it with an MSE loss, where, following the approach used in GEARS and scGPT, we randomly assign each perturbed cell with a control cell. The control cells are then perturbed by the model as described in Sec. 2, and the MSE between the predicted and the observed perturbed cells is calculated, also with higher weights on the top 20 DEGs (Eqs. 9 and 10). This ablated model is referred to as the MSE model. A summary of the ablated models are provided in Supplementary Table S2. All ablated models are evaluated following the procedure described in Sec. 3.2. The ΔPCC and W2 distance of ablated models on the Norman et al. dataset over 10 data splits are given in Fig. 3. Results on the other datasets and metrics are given in Supplementary Figs. S4, S5, and S6.

**Fig. 3.**
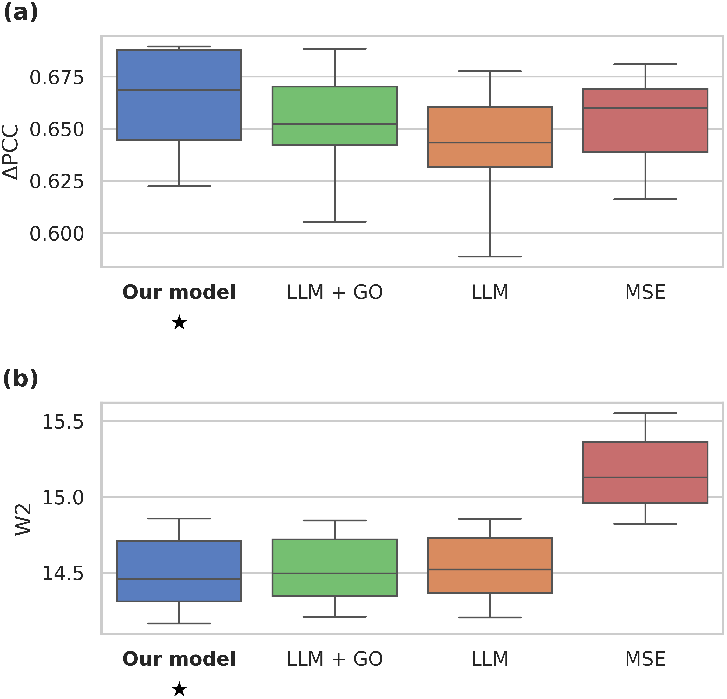
Ablation results on the Norman et al. dataset, evaluated using (a) Δ PCC and (b) W2 distance. The model with the best average result on each metric is bolded and starred.

Overall, our proposed model with all three gene representation components (LLM, GO, and STRING) and an OT-based loss consistently demonstrates strong performance relatively to ablation baselines, achieving either the best or the second-best result on all datasets and metrics. Model performance generally declines as gene representations are progressively removed, demonstrating the effectiveness of incorporating representations from multiple knowledge sources, consistent with the findings of Wenkel et al. [5]. The only exception is on the Adamson et al. dataset, where the LLM + GO variant achieves higher average ΔPCC and lower average W2 distance on DEGs than our proposed full model, while our model still achieves better average performance when considering all genes. Moreover, all models trained with an OT-based loss, including our proposed full model, the LLM + GO model, and the LLM model outperform the MSE variant in terms of the W2 distance, both on DEGs and on all genes. This result further validates the benefit of using an OT-based approach instead of randomly pairing up control and perturbed cells and not accounting for the inherent heterogeneity of scRNA-seq data.

### 3.4. Differential expression analysis

Next, following Wang et al. [8], we investigate the potential utility of our model in DE analysis. For this experiment, we first train the model on a single data split, and obtain the predictions on the test set. For each test perturbation, we perform DE analysis on both the predicted and the ground truth perturbed expression profiles using *t*-tests, which results in two sets of *t*-statistics. Since the degrees of freedom are fixed across genes, these *t*-statistics can be interpreted as DE scores, where genes with higher scores show stronger evidence of being up-regulated, and genes with lower scores show stronger evidence of being down-regulated. From the comparison between the control and the ground truth perturbed expression profiles, genes with adjusted *p*-values *<* 0.01 and log fold changes greater than 0 are considered the ground truth up-regulated, while those with adjusted *p*-values *<* 0.01 and log fold changes less than 0 are considered ground truth down-regulated [8]. Using the predicted DE scores and these ground truth gene sets, we calculate the area under the receiver operating characteristic curve (AUC) separately for the up- and down-regulated genes. As a summary statistic, we also compute the Spearman’s rank correlation between the predicted and the ground truth DE scores. We choose Spearman’s correlation since it is rank-based and only concerns whether the rankings of the genes are preserved, which aligns with the rank-based formulation of the AUC. The results across test perturbations on the Norman et al. dataset are summarized in Table 1. Results on other datasets are given in Supplementary Tables S3 and S4.

**Table 1.** Spearman’s correlation between predicted and ground truth DE scores, and AUCs for the classification of up- and down-regulated genes on the Norman et al. dataset. The best mean result for each metric is highlighted in bold.

| Model | Spearman’s corr. | AUC up-reg. | AUC down-reg. |
| --- | --- | --- | --- |
| Our model | **0.266 ± 0.097** | **0.793 ± 0.133** | 0.625 ± 0.069 |
| scLAMBDA | 0.225 ± 0.088 | 0.780 ± 0.130 | 0.623 ± 0.063 |
| scGPT | 0.066 ± 0.082 | 0.325 ± 0.087 | **0.787 ± 0.044** |
| GEARS | 0.160 ± 0.063 | 0.541 ± 0.106 | 0.583 ± 0.061 |

Overall, our model demonstrates a competitive performance on this DE analysis task, achieving either the best or the second-best average result across all datasets and metrics. The performance of scGPT shows a bias towards the identification of down-regulated genes, achieving the highest average AUC for down-regulated genes on both the Norman et al. and the Replogle et al. datasets. However, it achieves a low average AUC of 0.325 for up-regulated genes on the Norman et al. dataset, and thus the lowest average Spearman’s correlation of 0.066. On the other hand, our model achieves the highest average Spearman’s correlation on both the Norman et al. and the Replogle et al. dataset, showing its ability to most accurately recover the rankings of both up- and down-regulated genes. On the Adamson et al. dataset, scLAMDBA achieves the best Spearman’s correlation, as well as the highest AUCs on up- and down-regulated genes.

### 3.5. Genetic interaction modeling

Finally, we illustrate the biological relevance of our model by using its predictions on the Norman et al. dataset to compute genetic interaction (GI) scores for pairs of perturbed genes. Roohani et al. [1] considered various types of GIs in the Norman et al. dataset, including synergy, suppression, redundancy, neomorphism, and epistasis. These GI types were defined based on four GI metrics derived from perturbation response data: magnitude, similarity of single to double transcriptional profiles, model fit, and equality of contribution. For instance, a gene pair with sufficiently high magnitude score is considered synergistic, while one with low magnitude is classified as suppressive. For the derivation of these GI scores from perturbation data, please see Supplementary Sec. S6.

One potential application of computational methods in this context is to predict the outcomes of unseen double-gene perturbations and use this prediction to calculate the GI scores, thereby identifying potential interactions between the two genes. Following Roohani et al. [1] and Wang et al. [8], we assess models on this GI modeling task. Specifically, we randomly split the dataset into training, validation, and test sets, such that the test set only contains double-gene perturbations (see Supplementary Sec. S2). We then calculate the GI scores for each test perturbation using the predicted and the ground truth perturbed cells, and compute the MSEs between the two sets of scores across all test perturbations. This procedure is repeated 10 times for 10 random data splits. The average MSEs of all models on the four GI metrics are summarized in Table 2. For additional information on the implementation of this experiment, please see Supplementary Sec. S6.

**Table 2.** MSEs between the predicted and ground truth GI metrics on the Norman et al. dataset. The best mean MSE for each type of GI score is highlighted in bold.

| Model | Magnitude | Similarity | Model fit | Equal. of contrib. |
| --- | --- | --- | --- | --- |
| Our model | **0.111 ± 0.023** | **0.009 ± 0.004** | 0.007 ± 0.002 | **0.021 ± 0.006** |
| scLAMBDA | 0.194 ± 0.041 | 0.012 ± 0.003 | **0.006 ± 0.002** | 0.033 ± 0.008 |
| scGPT | 0.326 ± 0.088 | 0.024 ± 0.010 | 0.053 ± 0.019 | 0.031 ± 0.004 |
| GEARS | 0.339 ± 0.042 | 0.038 ± 0.008 | 0.049 ± 0.013 | 0.038 ± 0.008 |

Our model achieves the lowest average MSEs on all GI metrics, except on model fit, where scLAMBDA achieves the best average MSE of 0.006, followed closely by our model at 0.007. This result shows our model’s ability to capture biologically meaningful relationships between genes, demonstrating its potential as a tool to explore the genetic interaction landscape.

## 4. Discussion

In this work, we present an OT-based deep generative model that integrates multiple sources of prior knowledge to predict single-cell transcriptional responses to novel genetic perturbations. Across three benchmark datasets and four evaluation metrics, our model consistently demonstrates competitive performance with state-of-the-art methods. Ablation results further show that performance generally declines as gene representations from different knowledge sources are progressively dropped out. In addition, replacing the OT-based loss with an MSE loss on randomly paired predicted and observed perturbed cells degrades performance on population-wise similarity metrics, indicating suboptimal alignment between the predicted and observed perturbed cell distributions. These results suggest the effectiveness of the core components in our method design. Additional evaluations on differential expression analysis and genetic interaction modeling demonstrate the biological relevance of our model and suggest its potential applicability across a variety of contexts.

However, we acknowledge certain limitations of the proposed model. One limitation of our as well as other similar models is that they rely on external sources of gene information, so model performance might depend on the quality of these resources. In addition, in datasets where there are very few cells per perturbation, the added benefits of using an OT-based approach might diminish. For instance, in the extreme case where there is only one perturbed cell available, the OT loss reduces to an MSE between the expression of the perturbed cell and the average expression of the control cells. Additional experiments are needed to evaluate the performance of our model under such conditions.

Our model can be straightforwardly extended to predict responses to chemical perturbations or treatments by replacing the gene representations with chemical features, possibly from multiple knowledge sources. Even though our model is currently formulated for individual cell lines, it can be adapted to work on mixtures of cell types or samples from different donors – for example, by extending the encoder and decoder to additionally condition on cell-level and sample-level metadata. Another promising future direction is to incorporate information on the type of perturbation applied on each gene (e.g., knock-out, inhibition, activation), allowing the model to handle scenarios where multiple types of perturbation are present.

## Data availability

Our source code is available at https://github.com/minhchaudo/ot-gene-pert. All datasets used in this study are publicly available online at their respective repositories [11, 12, 13].

## Acknowledgements

We would like to acknowledge the computational resources provided by the Aalto Science-IT.

## References

1. Yusuf Roohani, Kexin Huang, and Jure Leskovec. Predicting transcriptional outcomes of novel multigene perturbations with gears. Nature Biotechnology, 42(6):927–935, June 2024. doi: 10.1038/s41587-023-01905-6.

2. Mohammad Lotfollahi, F. Alexander Wolf, and Fabian J. Theis. scgen predicts single-cell perturbation responses. Nature Methods, 16(8):715–721, August 2019. doi: 10.1038/s41592-019-0494-8.

3. Mohammad Lotfollahi, Anna Klimovskaia Susmelj, Carlo De Donno, et al. Predicting cellular responses to complex perturbations in high-throughput screens. Molecular Systems Biology, 19(6):e11517, June 2023. doi: 10.15252/msb.202211517.

4. Charlotte Bunne, Stefan G. Stark, Gabriele Gut, et al. Learning single-cell perturbation responses using neural optimal transport. Nature Methods, 20(11):1759–1768, November 2023. doi: 10.1038/s41592-023-01969-x.

5. Frederik Wenkel, Wilson Tu, Cassandra Masschelein, et al. Txpert: Leveraging biochemical relationships for out-of-distribution transcriptomic perturbation prediction. arXiv, 2025. doi: 10.48550/ARXIV.2505.14919.

6. Ding Bai, Caleb N Ellington, Shentong Mo, et al. Attentionpert: accurately modeling multiplexed genetic perturbations with multi-scale effects. Bioinformatics, 40(Supplement 1):i453–i461, June 2024. doi: 10.1093/bioinformatics/btae244.

7. Yiqun Chen and James Zou. Genepert: Leveraging genept embeddings for gene perturbation prediction. bioRxiv, October 2024. doi: 10.1101/2024.10.27.620513.

8. Gefei Wang, Tianyu Liu, Jia Zhao, et al. Modeling and predicting single-cell multi-gene perturbation responses with sclambda. bioRxiv, December 2024. doi: 10.1101/2024.12.04.626878.

9. Xiaoning Qi, Lianhe Zhao, Chenyu Tian, et al. Predicting transcriptional responses to novel chemical perturbations using deep generative model for drug discovery. Nature Communications, 15(1):9256, October 2024. doi: 10.1038/s41467-024-53457-1.

10. Jean Feydy, Thibault Séjourné, François-Xavier Vialard, et al. Interpolating between optimal transport and mmd using sinkhorn divergences. In Proceedings of the Twenty-Second International Conference on Artificial Intelligence and Statistics, volume 89 of Proceedings of Machine Learning Research, pages 2681–2690. PMLR, 16–18 Apr 2019. URL https://proceedings.mlr.press/v89/feydy19a.html.

11. Britt Adamson, Thomas M. Norman, Marco Jost, et al. A multiplexed single-cell crispr screening platform enables systematic dissection of the unfolded protein response. Cell, 167(7):1867–1882.e21, December 2016. doi: 10.1016/j.cell.2016.11.048.

12. Joseph M. Replogle, Thomas M. Norman, Albert Xu, et al. Combinatorial single-cell crispr screens by direct guide rna capture and targeted sequencing. Nature Biotechnology, 38(8): 954–961, August 2020. doi: 10.1038/s41587-020-0470-y.

13. Thomas M. Norman, Max A. Horlbeck, Joseph M. Replogle, et al. Exploring genetic interaction manifolds constructed from rich single-cell phenotypes. Science, 365(6455):786–793, August 2019. doi: 10.1126/science.aax4438.

14. Haotian Cui, Chloe Wang, Hassaan Maan, et al. scgpt: toward building a foundation model for single-cell multi-omics using generative ai. Nature Methods, 21(8):1470–1480, August 2024. doi: 10.1038/s41592-024-02201-0.

15. Michael Ashburner, Catherine A. Ball, Judith A. Blake, et al. Gene ontology: tool for the unification of biology. Nature Genetics, 25(1):25–29, May 2000. doi: 10.1038/75556.

16. The Gene Ontology Consortium, Suzi A Aleksander, James Balhoff, et al. The gene ontology knowledgebase in 2023. GENETICS, 224(1):iyad031, May 2023. doi: 10.1093/genetics/iyad031.

17. National Center for Biotechnology Information. Gene. URL https://www.ncbi.nlm.nih.gov/gene/. Accessed 2025-11-14.

18. Yiqun Chen and James Zou. Genept: A simple but effective foundation model for genes and cells built from chatgpt. bioRxiv, October 2023. doi: 10.1101/2023.10.16.562533.

19. Damian Szklarczyk, Rebecca Kirsch, Mikaela Koutrouli, et al. The string database in 2023: protein–protein association networks and functional enrichment analyses for any sequenced genome of interest. Nucleic Acids Research, 51(D1):D638–D646, January 2023. doi: 10.1093/nar/gkac1000.

