## Supplementary Information for "Predicting single-cell responses to novel genetic perturbations with optimal transport"

Chau Do and Harri Lähdesmäki

August, 2025

#### 1 Summary of datasets

Table 1 provides a summary of the three datasets used in our experiments.

Table 1: Summary of datasets.

| Dataset name | Number of perturbations ($P$) | Average number of cells per perturbation ($n_p$) |
|---|---|---|
| Adamson et al. | Single-gene: 81 | Single-gene: 514 |
| Replogle et al. | Single-gene: 645 | Single-gene: 100 |
| Norman et al. | Single-gene: 102, Double-gene: 128 | Single-gene: 466, Double-gene: 269 |

#### 2 Analysis details

**Model training** Our model is trained end-to-end using the Adam optimizer with initial learning rate $10^{-3}$. The learning rate is scheduled to decay by a factor of 0.2 every 30 epochs, following Wang et al. [1]. The number of epochs is tuned using the validation set, with the maximum number of epochs set to 100. The pretraining phase consists of 100 epochs with a learning rate of $10^{-3}$. The latent dimensions of both the perturbation representation $\mathbf{p}$ and the cell representations $\mathbf{Z}$ are set to 64. The single-cell loss is implemented using `geomloss.SamplesLoss("sinkhorn")` with `p=2`. The training loss is configured using the following coefficients: $\lambda_{\text{DEG}} = 1$, $\lambda_1 = 0.3$, and $\lambda_2 = 0.01$. Other models are trained following their publicly available instructions and implementations.

**Data splitting** For the perturbation response prediction task and the differential expression analysis task, perturbations are split into training, validation, and test sets using GEARS's `DataSplitter` function [2], with `train_gene_set_size = 0.75` and `combo_seen2_train_frac = 0.75`. Specifically, 25% of all perturbed genes are used for testing, and the remaining 75% are used for training and validation. When the dataset contains both single-gene and double-gene perturbations, all single-gene and double-gene perturbations with at least one gene in the test gene set are included in the test perturbation set, along with 25% of double-gene perturbations with both genes in the training gene set. This enables additional evaluation on double-gene perturbations with both genes already seen in training.

For the genetic interaction modeling task, the Norman et al. [3] dataset is randomly split into training, validation, and test sets, such that the test set only contains 50% of all double-gene perturbations in the dataset. Data splitting is performed using function `numpy.random.choice`.

**Gene representations** The LLM gene embeddings are generated by Wang et al. [1] using the GPT-3.5 model. The Jaccard GO similarities between genes are derived by Roohani et al. [2]. The STRING v11.5 protein association data [4] are preprocessed by Wenkel et al. [5].

**Perturbation filtering** GEARS could not predict the outcomes of perturbations involving genes that are missing from its GO graph. Such perturbations are thus removed from the benchmark datasets.

Perturbations involving genes missing from the set of NCBI-based embeddings derived by Wang et al. [1] cannot be predicted by scLAMBDA and are thus removed from the benchmark datasets.

ScGPT requires that perturbed genes have their expression data included in the dataset to be able to predict their perturbation effects. The Norman et al. [3] and Adamson et al. [6] datasets, which are preprocessed by Roohani et al. [2], include the expression values of all perturbed genes along with the top 5000 HVGs, so scGPT can predict the outcomes of all perturbations in these two datasets. However, the Replogle et al. [7]

dataset, also preprocessed by Roohani et al. [2], only includes the expressions of the top 5000 HVGs. Perturbed genes not among the top 5000 HVGs are excluded from the expression profiles of cells, possibly because this dataset includes a relatively large number of perturbed genes. Perturbations of such genes cannot be modeled by scGPT and are thus removed from this dataset.

### 3 Perturbation response prediction

Results of the perturbation response prediction experiment on the Norman et al., Adamson et al., and Replogle et al. datasets are given in Fig. 1, Fig. 2, and Fig. 3, respectively. A description of the experiment can be found in Sec. 3.2 of the main text.

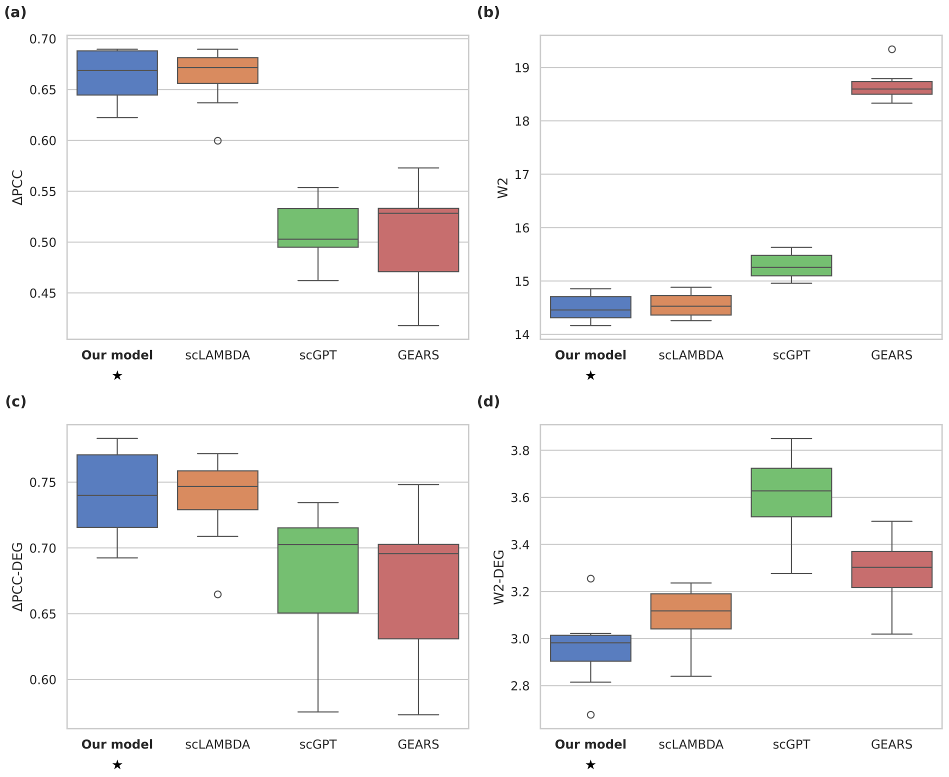


Figure 1: Performance of models on the Norman et al. dataset, evaluated using (a) $\Delta$ PCC, (b) W2 distance, (c) $\Delta$ PCC on top 20 DEGs, and (d) W2 distance on top 20 DEGs. The model with the best average result on each metric is bolded and starred.

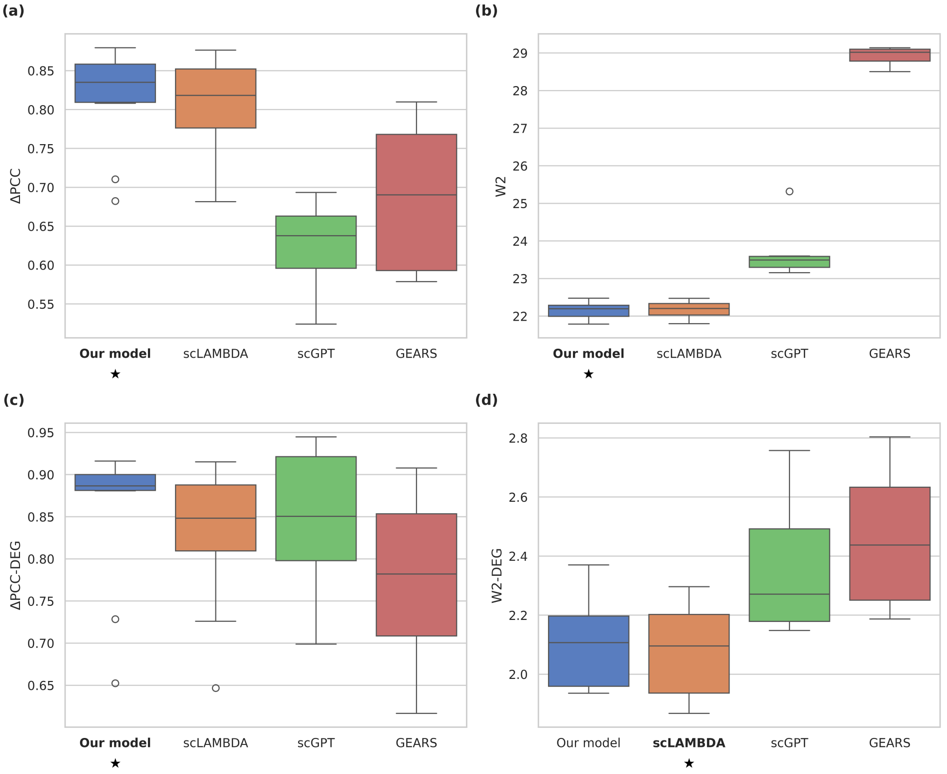


Figure 2: Performance of models on the Adamson et al. dataset, evaluated using (a) Δ PCC, (b) W2 distance, (c) Δ PCC on top 20 DEGs, and (d) W2 distance on top 20 DEGs. The model with the best average result on each metric is bolded and starred.

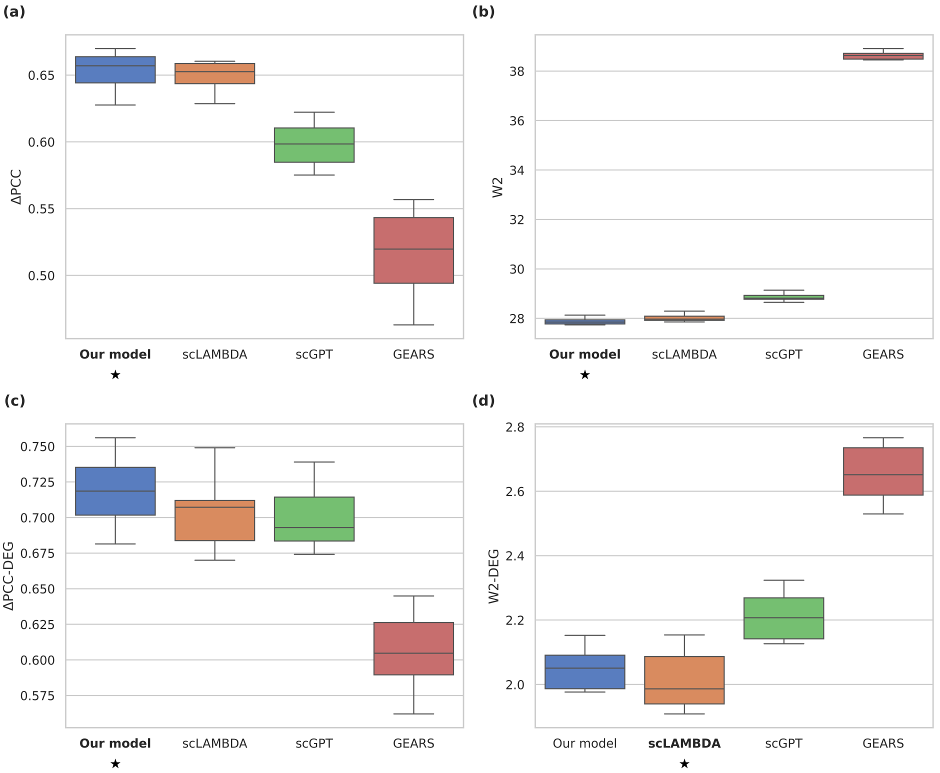


Figure 3: Performance of models on the Replogle et al. dataset, evaluated using (a) Δ PCC, (b) W2 distance, (c) Δ PCC on top 20 DEGs, and (d) W2 distance on top 20 DEGs. The model with the best average result on each metric is bolded and starred.

### 4 Ablation test

#### 4.1 Summary of ablated models

Table 2 provides a summary of the ablated models, which are described in Sec. 3.3 of the main text.

Table 2: Summary of ablated models. A check (✓) indicates the presence of a component, while the absence of a check indicates the absence of the corresponding component. In the MSE model, $\mathcal{L}_{sc}$ is replaced by an MSE term, also with higher weights on the top 20 DEGs as in $\mathcal{L}_{sc}$.

| Model | LLM | GO | STRING | $\mathcal{L}_{sc}$ |
|---|---|---|---|---|
| Full model | ✓ | ✓ | ✓ | ✓ |
| LLM + GO | ✓ | ✓ | | ✓ |
| LLM | ✓ | | | ✓ |
| MSE | ✓ | ✓ | ✓ | |

#### 4.2 Results

Ablation results on the Norman et al., Adamson et al., and Replogle et al. datasets are given in Fig. 4, Fig. 5, and Fig. 6, respectively. A description of the experiment can be found in Sec. 3.3 of the main text.

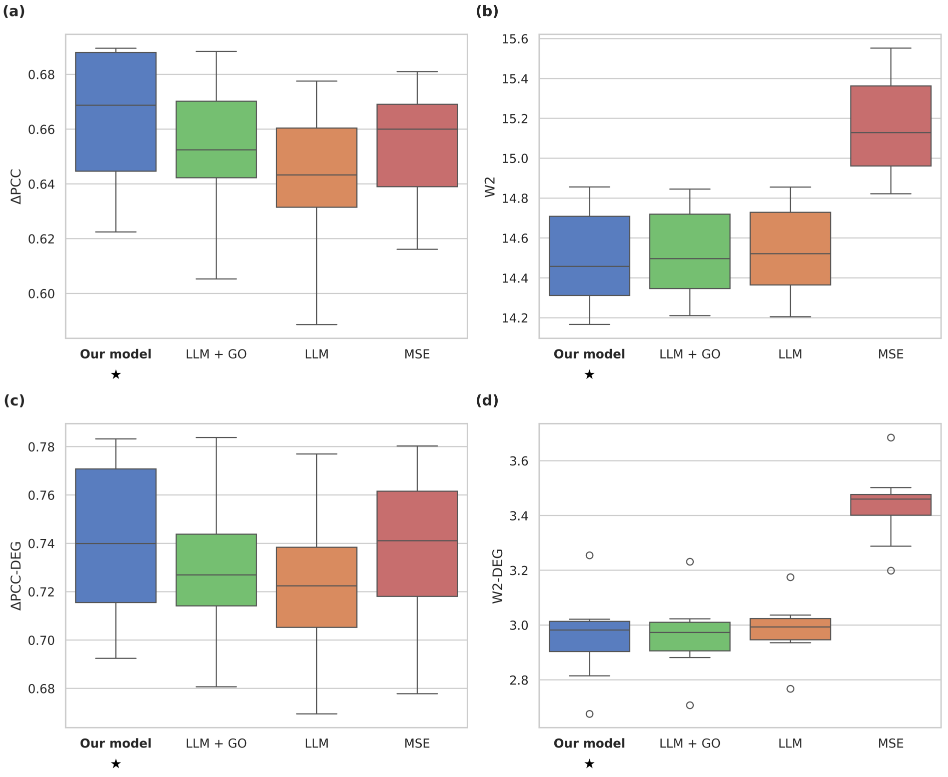


Figure 4: Performance of the full model and ablated models on the Norman et al. dataset, evaluated using (a) Δ PCC, (b) W2 distance, (c) Δ PCC on top 20 DEGs, and (d) W2 distance on top 20 DEGs. The model with the best average result on each metric is bolded and starred.

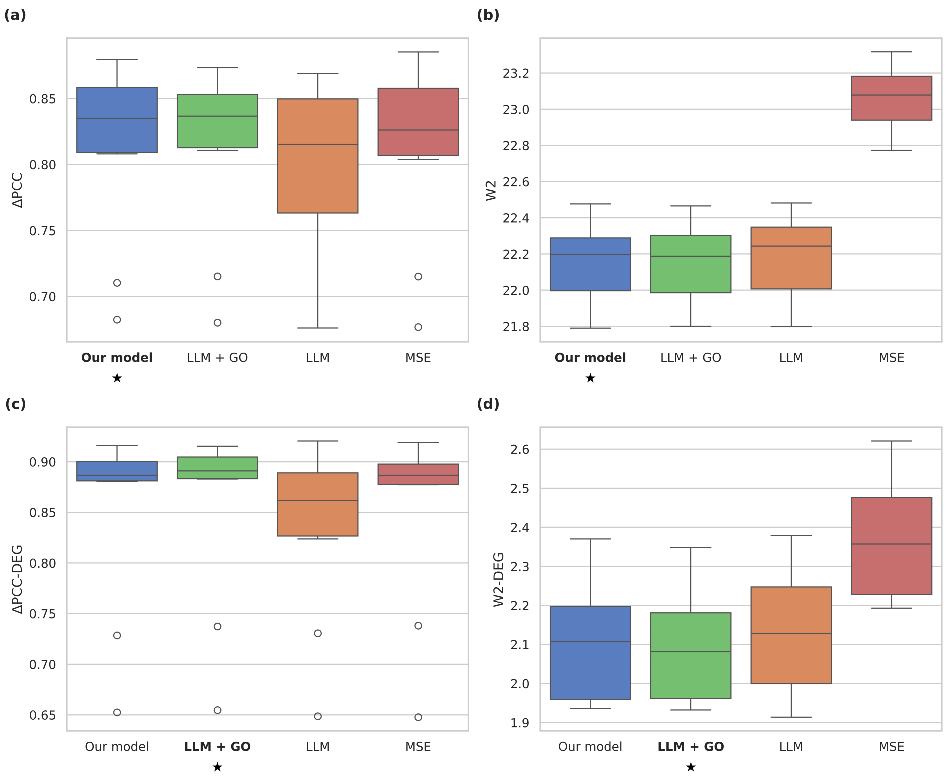


Figure 5: Performance of the full model and ablated models on the Adamson et al. dataset, evaluated using (a) $\Delta$ PCC, (b) W2 distance, (c) $\Delta$ PCC on top 20 DEGs, and (d) W2 distance on top 20 DEGs. The model with the best average result on each metric is bolded and starred.

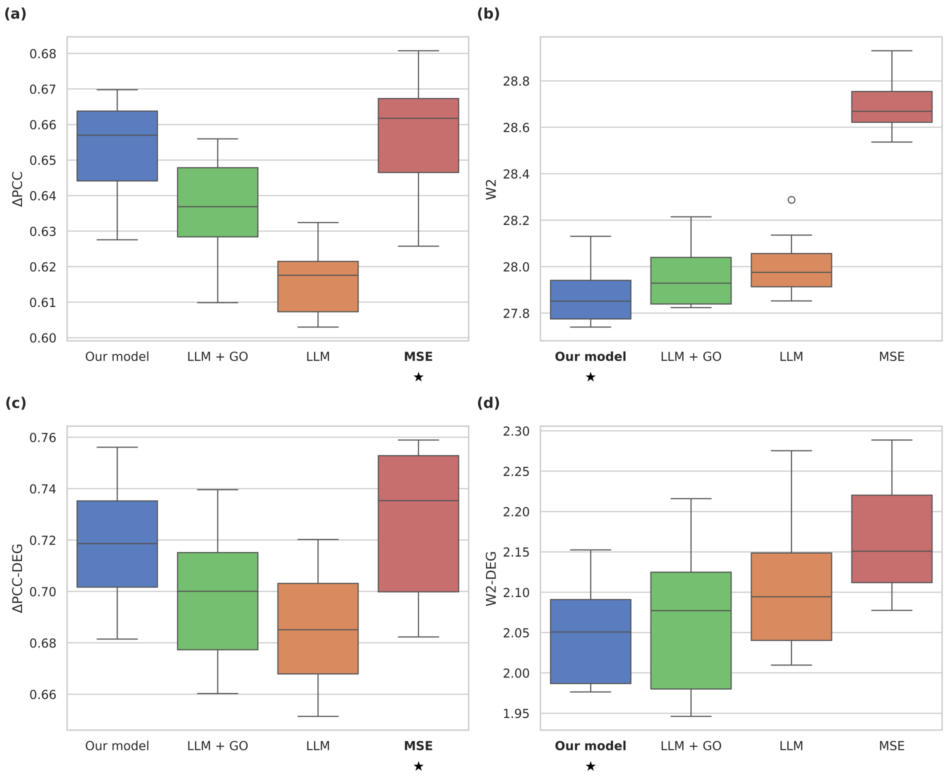


Figure 6: Performance of the original model and ablated models on the Replogle et al. dataset, evaluated using (a) $\Delta$ PCC, (b) W2 distance, (c) $\Delta$ PCC on top 20 DEGs, and (d) W2 distance on top 20 DEGs. The model with the best average result on each metric is bolded and starred.

#### 5 Differential expression analysis

Differential expression analysis results on the Adamson et al. and Replogle et al. datasets are given in Table 3 and Table 4, respectively. A description of this experiment can be found in Sec. 3.4 of the main text.

Table 3: Spearman's correlation between predicted and ground truth DE scores, and AUCs for the classification of up- and down-regulated genes on the Adamson et al. dataset. The best mean result for each metric is highlighted in bold.

| Model | Spearman's corr. | AUC up-reg. | AUC down-reg. |
|---|---|---|---|
| Our model | 0.659 ± 0.081 | 0.880 ± 0.045 | 0.770 ± 0.037 |
| scLAMBDA | **0.689 ± 0.080** | **0.896 ± 0.047** | **0.790 ± 0.032** |
| scGPT | 0.247 ± 0.063 | 0.701 ± 0.035 | 0.580 ± 0.050 |
| GEARS | 0.322 ± 0.051 | 0.609 ± 0.023 | 0.681 ± 0.048 |

Table 4: Spearman's correlation between predicted and ground truth DE scores, and AUCs for the classification of up- and down-regulated genes on the Replogle et al. dataset. The best mean result for each metric is highlighted in bold.

| Model | Spearman's corr. | AUC up-reg | AUC down-reg. |
|---|---|---|---|
| Our model | **0.528 ± 0.234** | 0.829 ± 0.138 | 0.790 ± 0.138 |
| scLAMBDA | 0.524 ± 0.240 | **0.837 ± 0.134** | 0.781 ± 0.154 |
| scGPT | 0.506 ± 0.223 | 0.785 ± 0.149 | **0.803 ± 0.132** |
| GEARS | 0.374 ± 0.189 | 0.726 ± 0.134 | 0.746 ± 0.121 |

#### 6 Genetic interaction modeling

Following Roohani et al. [2] and Norman et al. [3], the GI scores (metrics) are calculated by first fitting a linear model:

$$\delta\bar{\mathbf{y}}_{a,b} = c_a\delta\bar{\mathbf{y}}_a + c_b\delta\bar{\mathbf{y}}_b, \tag{1}$$

where $c_a$ and $c_b$ are model coefficients, $\delta\bar{\mathbf{y}}_a = \text{mean}(\mathbf{Y}_a) - \text{mean}(\mathbf{X})$ is the vector of pseudobulk expression changes compared to the control when gene $a$ is perturbed, and $\delta\bar{\mathbf{y}}_{a,b}$ is the vector of pseudobulk expression changes when both genes $a$ and $b$ are perturbed. This regression model is fitted using a Theil-Sen estimator on genes with average expressions greater than 0.5 unique molecule identifiers (UMIs) per cell.

Given this regression model, GI metrics are defined following the implementation of Roohani et al. [2] as follows:

- Magnitude: $\sqrt{c_a^2 + c_b^2}$
- Similarity of single to double transcriptional profiles: $\text{dcor}([\mathbf{a}, \mathbf{b}], \mathbf{ab})$
- Model fit: $\text{corr}(c_a\mathbf{a} + c_b\mathbf{b}, \mathbf{ab})$
- Equality of contribution: $\frac{\min(\text{dcor}(\mathbf{a},\mathbf{ab}),\text{dcor}(\mathbf{b},\mathbf{ab}))}{\max(\text{dcor}(\mathbf{a},\mathbf{ab}),\text{dcor}(\mathbf{b},\mathbf{ab}))}$

Here, $\mathbf{a}$, $\mathbf{b}$, and $\mathbf{ab}$ are abbreviations for $\delta\bar{\mathbf{y}}_a$, $\delta\bar{\mathbf{y}}_b$, and $\delta\bar{\mathbf{y}}_{a,b}$, respectively, $\text{dcor}(\cdot)$ denotes the distance correlation function, $\text{corr}(\cdot)$ denotes the Pearson correlation coefficient function, and $[\mathbf{a}, \mathbf{b}]$ denotes concatenation of two column vectors along the second dimension.

#### 7 Review of baselines

GEARS [2] is a GNN-based method that leverages a network of gene-gene relationships to derive embeddings for novel perturbed genes. Specifically, the gene network is derived from the GO database [8, 9], with each edge weight representing the Jaccard similarity between the two sets of pathway-related GO terms associated with the two genes. This design reflects the intuition that genes participating in similar pathways should have similar biological roles and thus have similar effects on expression profiles when perturbed.

ScLAMBDA [1] is a deep generative model with a variational autoencoder-based architecture. In scLAMBDA, novel perturbed genes are represented using LLM embeddings of gene descriptions retrieved from the NCBI Gene database, i.e., GenePT embeddings [10, 11]. During training, scLAMBDA inputs the expression profiles of perturbed cells and disentangles the basal cell state from the perturbation effect in the latent space. To ensure effective disentanglement, the mutual information between the basal cell state and the perturbation representation is additionally minimized. During test time evaluation, scLAMBDA combines the basal cell state with the perturbation representation to generate perturbed cells.

ScGPT [12] is a foundation model for single-cell multi-omics, leveraging a self-attention transformer architecture. Pretrained on large-scale single-cell sequencing datasets, scGPT can be adapted for different downstream analyses, including perturbation outcome prediction. For this task, scGPT additionally inputs a vector indicating the perturbation condition of each gene (i.e., perturbed or not perturbed), which influences the final embedding of the cell. As a result, scGPT requires that perturbed genes have their expression data included in the dataset in order to predict their perturbation effects. In our experiments, we initialize the model with parameters from scGPT's whole-human checkpoint, which was pretrained on 33 million cells, and fine-tune the model on each perturbation dataset.

TxPert [5] also explores the idea of integrating different sources of gene information, specifically by extending the GNN-based architecture of GEARS to support multiple knowledge graphs. However, TxPert is limited to graph-based prior knowledge, i.e., priors in the form of gene-gene relationships, while our model directly utilizes gene representation vectors and is thus capable of incorporating both gene-gene relationships and more general gene embeddings (e.g., LLM-based embeddings of gene descriptions). While TxPert is a newly released model with strong reported performance, the reported model uses proprietary data sources to derive gene-gene relationship information. The public release thus provides an ablated version that only uses public data sources (GO and STRING) to represent gene-gene relationships. Furthermore, the provided software currently only supports inference using the pretrained ablated model and does not provide an entry point for training from scratch that would be needed for fair performance comparisons. For these reasons, we exclude TxPert from our comparative analysis.

### References


[1] Gefei Wang, Tianyu Liu, Jia Zhao, et al. Modeling and predicting single-cell multi-gene perturbation responses with sclambda. *bioRxiv*, December 2024. doi: 10.1101/2024.12.04.626878.

[2] Yusuf Roohani, Kexin Huang, and Jure Leskovec. Predicting transcriptional outcomes of novel multi-gene perturbations with gears. *Nature Biotechnology*, 42(6):927–935, June 2024. doi: 10.1038/s41587-023-01905-6.

[3] Thomas M. Norman, Max A. Horlbeck, Joseph M. Replogle, et al. Exploring genetic interaction manifolds constructed from rich single-cell phenotypes. *Science*, 365(6455):786–793, August 2019. doi: 10.1126/science.aax4438.

[4] Damian Szklarczyk, Annika L Gable, David Lyon, et al. String v11: protein–protein association networks with increased coverage, supporting functional discovery in genome-wide experimental datasets. *Nucleic Acids Research*, 47(D1):D607–D613, January 2019. doi: 10.1093/nar/gky1131.

[5] Frederik Wenkel, Wilson Tu, Cassandra Masschelein, et al. Txpert: Leveraging biochemical relationships for out-of-distribution transcriptomic perturbation prediction. *arXiv*, 2025. doi: 10.48550/ARXIV.2505.14919.

[6] Britt Adamson, Thomas M. Norman, Marco Jost, et al. A multiplexed single-cell crispr screening platform enables systematic dissection of the unfolded protein response. *Cell*, 167(7):1867–1882.e21, December 2016. doi: 10.1016/j.cell.2016.11.048.

[7] Joseph M. Replogle, Thomas M. Norman, Albert Xu, et al. Combinatorial single-cell crispr screens by direct guide rna capture and targeted sequencing. *Nature Biotechnology*, 38(8):954–961, August 2020. doi: 10.1038/s41587-020-0470-y.

[8] Michael Ashburner, Catherine A. Ball, Judith A. Blake, et al. Gene ontology: tool for the unification of biology. *Nature Genetics*, 25(1):25–29, May 2000. doi: 10.1038/75556.

[9] The Gene Ontology Consortium, Suzi A Aleksander, James Balhoff, et al. The gene ontology knowledgebase in 2023. *GENETICS*, 224(1):iyad031, May 2023. doi: 10.1093/genetics/iyad031.

[10] National Center for Biotechnology Information. Gene. URL `https://www.ncbi.nlm.nih.gov/gene/`. Accessed 2025-11-14.

[11] Yiqun Chen and James Zou. Genept: A simple but effective foundation model for genes and cells built from chatgpt. *bioRxiv*, October 2023. doi: 10.1101/2023.10.16.562533.

[12] Haotian Cui, Chloe Wang, Hassaan Maan, et al. scgpt: toward building a foundation model for single-cell multi-omics using generative ai. *Nature Methods*, 21(8):1470–1480, August 2024. doi: 10.1038/s41592-024-02201-0.
